# Indole-3-glycerolphosphate synthase, a branchpoint for the biosynthesis of tryptophan, indole, and benzoxazinoids in maize

**DOI:** 10.1101/2021.01.04.425338

**Authors:** Annett Richter, Adrian F. Powell, Mahdieh Mirzaei, Lucy J. Wang, Navid Movahed, Julia K. Miller, Miguel A. Piñeros, Georg Jander

## Abstract

The maize (*Zea mays*) genome encodes three indole-3-glycerolphosphate synthase enzymes (IGPS1, 2, and 3) catalyzing the conversion of 1-(2-carboxyphenylamino)-l-deoxyribulose-5-phosphate to indole-3-glycerolphosphate. Three further maize enzymes (BX1, benzoxazinoneless 1; TSA, tryptophan synthase α subunit; and IGL, indole glycerolphosphate lyase) convert indole-3-glycerolphosphate to indole, which is released as a volatile defense signaling compound and also serves as a precursor for the biosynthesis of tryptophan and defense-related benzoxazinoids. Phylogenetic analyses showed that IGPS2 is similar to enzymes found in both monocots and dicots, whereas maize IGPS1 and IGPS3 are in monocot-specific clades. Fusions of yellow fluorescent protein (YFP) with maize IGPS enzymes and indole-3-glycerolphosphate lyases were all localized in chloroplasts. In bimolecular fluorescence complementation assays, IGPS1 interacted strongly with BX1 and IGL, IGPS2 interacted primarily with TSA, and IGPS3 interacted equally with all three indole-3-glycerolphosphate lyases. Whereas *IGPS1* and *IGPS3* expression was induced by insect feeding, *IGPS2* expression was not. Transposon insertions in *IGPS1* and *IGPS3* reduced the abundance of both benzoxazinoids and free indole. *Spodoptera exigua* (beet armyworm) larvae show improved growth on *igps1* mutant maize plants. Together, these results suggest that IGPS1 and IGPS3 function mainly in the biosynthesis of defensive metabolites, whereas IGPS2 may be involved in the biosynthesis of tryptophan. This metabolic channeling is similar, though less exclusive than that proposed for the three maize indole-3-glycerolphosphate lyases.

## Introduction

Grasses, including maize (*Zea mays*), produce a wide variety of constitutive and inducible defense compounds to protect themselves against herbivores and pathogens. Benzoxazinoids, a class of indole-derived antifeedant and insecticidal compounds in maize, provide resistance to numerous arthropod pests (McMullen *et al*., 2009a, Meihls *et al*., 2012, Meihls *et al*., 2013, Tzin *et al*., 2017, Bui *et al*., 2018). The core benzoxazinoid biosynthesis pathway is well-studied and involves seven BX enzymes (BX1-BX5, BX8, and BX9) that catalyze the formation of 2,4-dihydroxy-1,4-benzoxazin-3-one glucoside (DIBOA-Glc) from indole-3-glycerol phosphate (IGP) (Figure 1; Frey *et al*., 1997, Niemeyer, 2009, Wouters *et al*., 2016). The first reaction in the pathway is catalyzed by BX1, an indole-producing indole-3-glycerolphosphate lyase. Based on global coexpression network analysis, Wisecaver *et al*. (2017) proposed that the committed step in the benzoxazinoid biosynthesis pathway may be upstream of BX1. *IGPS1* (GRMZM2G106950), a maize gene that is co-regulated with BX genes in maize inbred line B73, was predicted to encode indole-3-glycerolphosphate synthase (IGPS), which catalyzes the ring closure of 1-(2-carboxyphenylamino)-l-deoxyribulose-5-phosphate into IGP (Figure 1). Maize contains two additional predicted IGPS genes, *IGPS2* (GRMZM2G169516) and *IGPS3* (GRMZM2G145870), but the expression of these genes was not co-regulated with benzoxazinoid pathway genes (Wisecaver *et al*., 2017).

**Figure 1.**
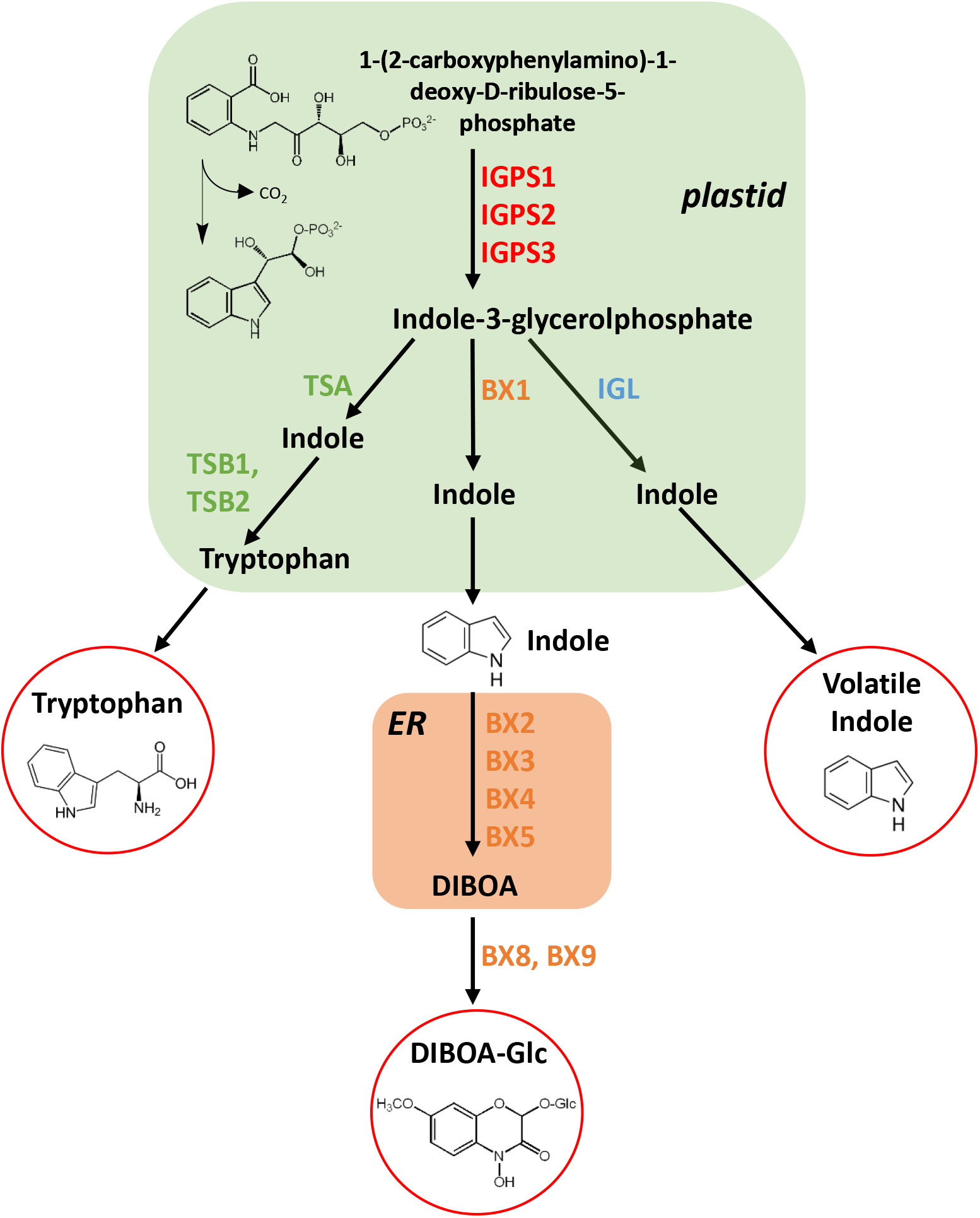
Biosynthetic pathways from indole-3-glycerolphosphate to DIBOA-Glc, tryptophan, and free indole in maize. Protein localization and confirmation of IGPS1, IPGS2, and IGPS3 enzymatic activity is described in this manuscript. IGPS = indole-3-glycerolphosphate synthase, BX = benzoxazinoneless, TSA = tryptophan synthase alpha subunit, TSB = tryptophan synthase beta subunit, IGL = indole-3-glycerolphosphate lyase, DIBOA-Glc = 2,4-dihydroxy-1,4-benzoxazin-3-one glucoside, ER = endoplasmic reticulum.

To date, the only plant IGPS enzymes that have been characterized are AT2G04400 and AT5G48220 in *Arabidopsis thaliana* (Arabidopsis; Li *et al*., 1995a). AT2G04400 enzymatic activity was verified by complementing an *Escherichia coli* mutant strain containing a missense mutation in the *trpC* gene, which encodes a bifunctional IGPS/*N*-(5’-pliosplioribosvi)-anthranilate isomerase (PRAI) enzyme (Yanofsky *et al*., 1971, Li *et al*., 1995a). Based on the complementation assays using *E. coli* mutants deficient in either the IGPS or the PRAI activity of TrpC, it appears that plants only have monofunctional IGPS enzymes (Li *et al*., 1995a, Li *et al*., 1995b). Indole-3-glycerolphosphate synthase activity was also verified in a wheat tissue extract, though the actual enzyme was not functionally characterized (Singh and Widholm, 1974).

Indole-3-glycerol phosphate (IGP) is a precursor for indole and numerous other indole-derived plant metabolites, including tryptophan, indole alkaloids, glucosinolates, benzoxazinoids, and auxin (Rose and Last, 1994, Frey *et al*., 2009). Arabidopsis has two confirmed indole-3-glycerolphosphate lyases, TSA1 (tryptophan synthase alpha subunit; AT3G54640) and INS (indole synthase; AT4G02610), which convert IGP to indole. Whereas TSA1 is involved in the tryptophan biosynthesis, as part of a complex with TSB1 (tryptophan synthase beta subunit AT5G54810), INS catalyzes indole synthesis independent of TSB (Zhang *et al*., 2008). Unlike TSA1, Arabidopsis INS does not have a chloroplast targeting sequence. Thus, the pathways for the biosynthesis of tryptophan and tryptophan-independent indole-containing metabolites may be separated in Arabidopsis subcellular compartments.

Three maize genes, *BX1 (benzoxazinoneless 1*), *TSA* (*tryptophan synthase α subunit*) and *IGL* (*indole glycerolphosphate lyase*), encode enzymes that convert IGP to indole (Figure 1; Kriechbaumer *et al*., 2008). TSA-GFP fusion proteins were chloroplast-localized in maize protoplast expression expriements (Kriechbaumer *et al*., 2008) and assays with spinach chloroplasts expressing maize *BX1* also indicated a chloroplastic location for this protein (Stettner, 1998). A fourth protein, designated as TSA-like, was localized to the cytosol and did not have indole biosynthesis activity *in vitro* (Kriechbaumer *et al*., 2008). *BX1, TSA*, and *IGL* genes can be differentiated by both their expression levels and their relative contributions to the biosynthesis of tryptophan, benzoxazinoids, or free indole. Although *BX1* is constitutively expressed in young maize seedlings, its expression also is induced by caterpillar feeding (Maag *et al*., 2016, Tzin *et al*., 2017). Similarly, *IGL* expression is upregulated in response to herbivory, consistent with the observed herbivory-induced increase in maize indole emission (Frey *et al*., 2000). Mutations in *IGL*, but not *BX1*, led to an indole-deficient phenotype (Ahmad *et al*., 2011). Conversely, only very low amounts of benzoxazinoids can be detected in *bx1* mutant maize lines (Hamilton, 1964, Melanson *et al*., 1997, Betsiashvili *et al*., 2015, Tzin *et al*., 2017), indicating that the indole produced by IGL and/or TSA is not easily incorporated into benzoxazinoids. Nevertheless, exogenous indole application to *bx1* mutant maize plants leads to complementation of the benzoxazinoid-deficient phenotype (Frey *et al*., 1997, Ahmad *et al*., 2011).

Maize TSA forms a complex with TSB1 *in vitro*. Partial purification and proteomic analysis of the 160 kD tryptophan synthase complex from maize leaves identified TSA and TSB proteins, but not BX1, IGL, or TSA-like (Kriechbaumer *et al*., 2008). To date, no maize *TSA* mutation has been reported, but double mutations in the tryptophan synthase ß-subunit genes *TSB1* and *TSB2* led to a seedling-lethal phenotype with high indole accumulation (Wright *et al*., 1992). Maize plants with double mutations in *BX1* and *IGL1* are viable, indicating that neither of these enzymes makes an essential contribution to tryptophan biosynthesis (Ahmad *et al*., 2011)

Similar to IGP catabolism by the *BX1, TSA*, and *IGL* protein products, IGP synthesis was predicted to be encoded by three maize genes, *IGPS1*, *IGPS2*, and *IGPS3*. Here we report experiments to confirm the function of maize IGPS enzymes and determine whether they represent an earlier branch point in the indole biosynthesis pathway. Bimolecular fluorescence complementation (BIFC) assays show differences in the interaction profiles. For instance, whereas IGPS1 strongly interacts with BX1 and IGL but not with TSA, IGPS2 has a stronger interaction with TSA. However, the analysis of *IGPS1* and *IGPS3* mutant plants suggests that the metabolic channeling of IGP is not as strong as that of indole produced by BX1, TSA, and IGL.

## Results

### Maize has three indole-3-glycerol phosphate synthase genes

Amino acid sequence comparisons showed greater identity between maize IGPS1 and IGPS3 (64%) than when comparing IGPS1 *vs*. IGPS2 (52%) and IGPS3 *vs*. IGPS2 (51 %). We performed a phylogenetic analysis to investigate the evolutionary relationships of the IGPS family in monocots and dicots, including maize, Arabidopsis, *Triticum aestivum* (wheat) *Sorghum bicolor* (sorghum), *Oryza sativa* (rice), *Hordeum vulgare* (barley), *Setaria viridis* (green bristlegrass), *Populus trichocarpa* (poplar), and *Nicotiana attenuata* (coyote tobacco). Figure 2). Maize IGPS2 was in a cluster with both monocot and dicot enzymes, including the two Arabidopsis enzymes with confirmed IGP synthesis activity (Zhang *et al*., 2008). By contrast, IGPS1 and IGPS3 were in separate clades that contain only monocot enzymes. Phylogenetically, the maize IGPS enzymes were most similar to those of a close relative, *S. bicolor* (Springer et al., 1989).

**Figure 2.**
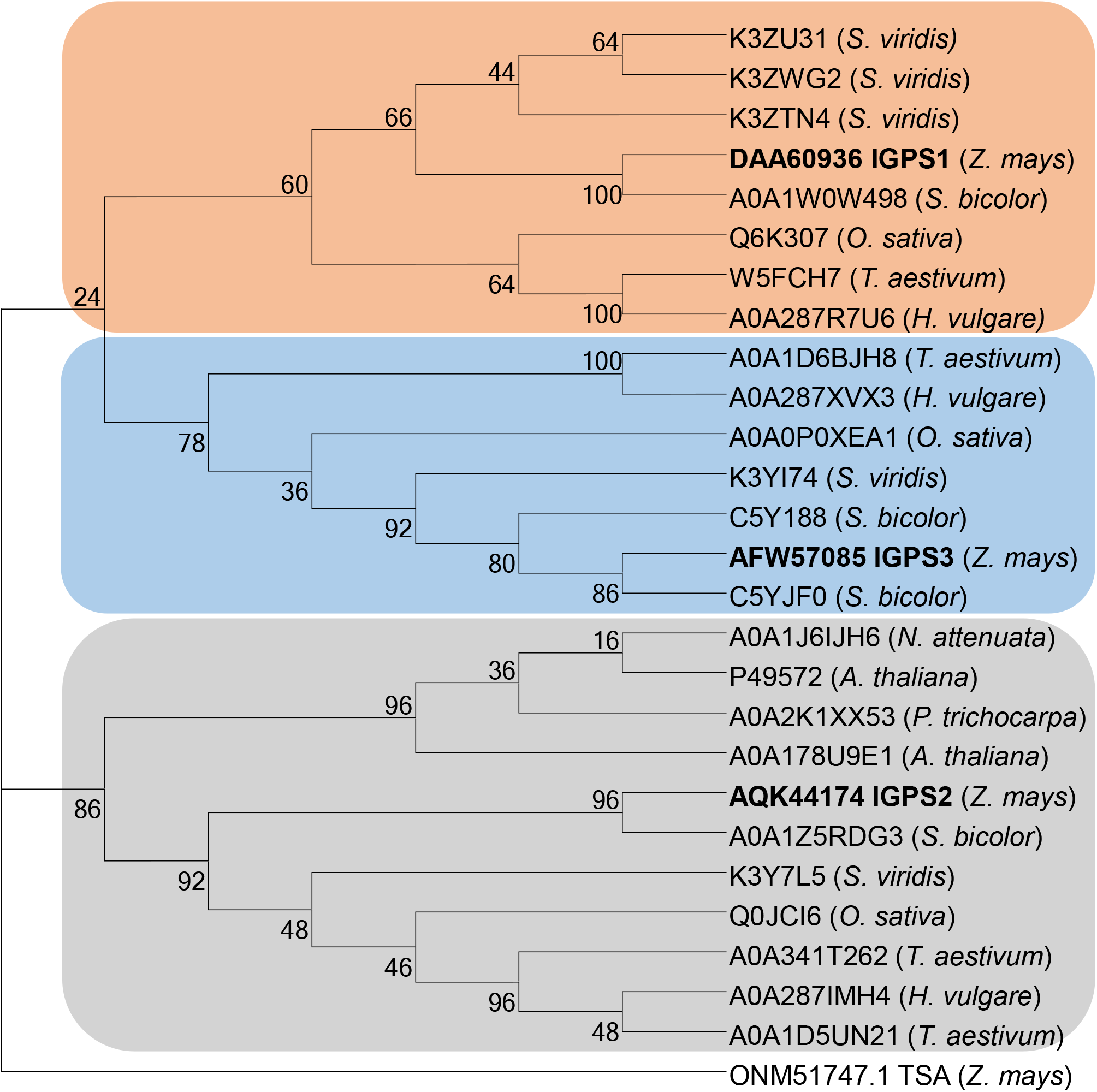
Phylogenetic analysis of indole-3-glycerol synthase (IGPS) proteins from monocots and dicots, including *Zea mays* (maize), *Triticum aestivum* (wheat) *Sorghum bicolor* (sorghum), *Oryza sativa* (rice), *Hordeum vulgare* (barley), *Setaria viridis* (green bristlegrass), *Arabidopsis thaliana* (mouse-ear cress), *Populus trichocarpa* (poplar), and *Nicotiana attenuata* (coyote tobacco). The enzymes cluster into phylogenetic groups that include the three maize enzymes, IGPS1 (orange cluster); IGPS2 (grey cluster); and IGPS3 (blue cluster). The MUSCLE Codon algorithm was used to perform a sequence alignment, which was used to reconstruct the tree using the Maximum Likelihood method and JTT matrix-based model. The percentage of trees in which the associated taxa clustered together is shown next to the branches (1,000 bootstrap trials). Maize tryptophan synthase alpha subunit (TSA) was used as an outgroup for the analysis. All protein sequences were retrieved from GenBank (https://www.ncbi.nlm.nih.gov/genbank/) using the indicated identifiers.

Aligning the B73 alleles of the three *IGPS* genes with the genomes of the 26 founder lines of the maize Nested Association Mapping (NAM) population (McMullen *et al*., 2009b; www.gramene.org) showed high sequence conservation among members of this enzyme family, with 96-100% identity in the predicted protein sequence (Supplementary Table S2). The only exception was inbred line M162W, where *IGPS2* was partially aligned to two adjacent genes, which may result from an incorrect annotation of the M162W genome. The chromosome positions of the three *IGPS* genes were conserved in the 26 maize genome assemblies, with the inbred line Tzi8 being the exception, having a second, almost-identical, *IGPS1* gene in an assembled scaffold that was not associated with a chromosome. This is possibly an assembly error, as none of the other NAM founder lines genomes had any additional predicted *IGPS* genes.

The IGPS enzyme family is known for catalyzing a carbon-carbon ring-forming reaction in which the carbon atoms become covalently bonded (Wierenga, 2001). The three maize IGPS proteins have the characteristic domains of this enzyme class, including a TIM (triosephosphate isomerase)-like beta/alpha barrel domain, which is one of the most common enzyme folds (Supplemental Figure S1; Chan et al., 2017; Nagano et al., 2002). The TIM barrel motif includes a phosphate-binding domain, which is a conserved protein fold consisting of eight α-helices and eight parallel β-strands that alternate along the peptide backbone (Farber and Petsko, 1990, Brändén, 1991, Wilmanns *et al*., 1991). The well-organized scaffold contains predicted binding sites for indole, phosphate, ribulose/trilose, and 1-(2-carboxyphenylamino)-1-deoxy-D-ribulose-5-phosphate (Supplemental Figure S1).

To confirm the enzymatic activity of IGPS1, IGPS2, and IGPS3, we complemented the tryptophan-auxotrophic phenotype of the *E. coli trpC9800*^-^ strain (Yanofsky *et al*., 1971), which is deficient in the IGPS but not the PRAI enzymatic activity of TrpC. The coding regions of the three maize *IGPS* genes, including the signal sequences, were cloned into *E. coli* plasmid vectors for expression in *E. coli trpC9800*. Whereas the empty vector control strain grew only when the minimal medium was supplemented with tryptophan, all three maize *IGPS* genes complemented the tryptophan-auxotrophic phenotype (Figure 3).

**Figure 3.**
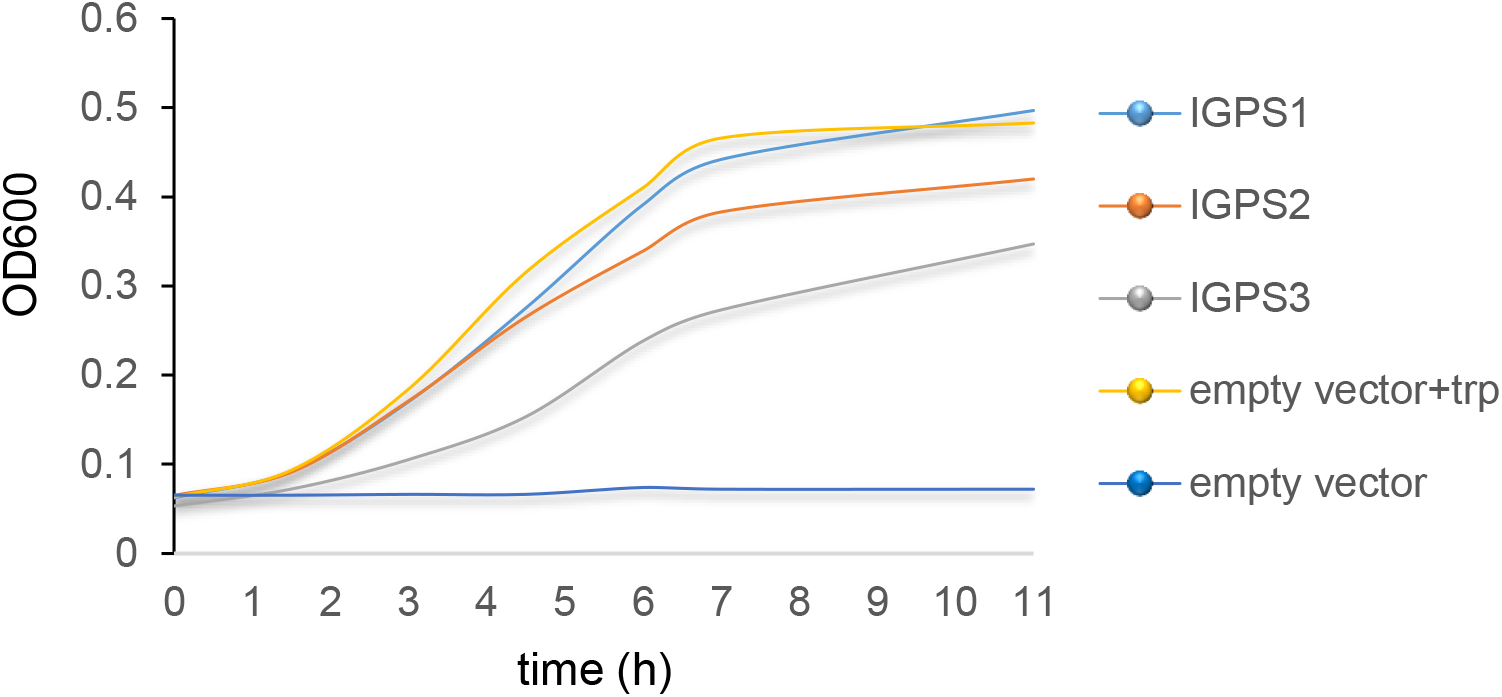
Identification of maize indole-3-glycerol-phosphate synthase (IGPS) activity. Maize IGPS enzyme activity was verified by cloning the cDNA and complementing an *E. coli trpC9800*^-^ mutation, which causes tryptophan auxotrophy through loss of indole-3-glycerolphosphate synthase activity. Empty vector controls were grown with or without the addition of 20 μg/ml tryptophan to the minimal medium. Optical density at 600 nm (OD600) was measured over 11 h.

### Maize *IGPS1* and *IGPS3* expression is induced by herbivory

Previous experiments showed that 24 h of *S. exigua* caterpillar feeding on maize inbred line B73 resulted in a 100-fold induction of *IGPS1* expression (p = < 0.001, Dunnett’s test) and a 5-fold induction of *IGPS3* (p = < 0.001, Dunnett’s test), but no significant changes in *IGPS2* (Tzin *et al*., 2017, Wisecaver *et al*., 2017; Supplemental Figure S2A). To determine whether the gene expression response is similar in the W22 inbred line, we infested maize seedlings with *S. exigua* for 24 h and measured the transcript levels by quantitative real-time PCR (qRT-PCR). Similar to the previous results with inbred line B73 (Supplemental Figure S2A), this showed significant induction of *IGPS1* (p < 0.001, *t-*test) and *IGPS3* (p < 0.001, *t-*test) transcripts (Figure 4A) and no significant transcriptional changes for *IGPS2*. Methyl jasmonate treatment, which regulates many insect defense responses in plants (Howe and Jander 2008; Martin et al., 2002; Richter et al., 2015), also increased *IGPS1* and *IGPS3* but not *IGPS2* expression (Figure 4B). We measured *IGPS* expression by 3’RNAseq (Kremling *et al*., 2018) in a time course with 0 h, 2 h, 8 h, and 24 h of *R. maidis* feeding. *IGPS1* expression showed a significant increase after 8 h of aphid feeding W22 (p < 0.05, *t-*test; Figure 4C), which is comparable to the significant induction at 4 h after *R. maidis* infestation in inbred line B73 (Tzin *et al*., 2015; Figure S2B). Although *IGPS3* expression was not significantly changed in response to *R. maidis* feeding, the Illumina 3’RNAseq read counts of *IGPS3* (in average: 188) in W22 plants were higher than those of *IGPS1* (average: 2.5) and *IGPS2* (average: 21), even in the absence of insect feeding. These results are different from the 3’RNAseq results with inbred line B73, which showed an average of 30 read counts for *IGPS1*, 18 read counts for *IGPS3*, and 12 read counts for *IGPS2* (Tzin *et al*., 2015). Given that benzoxazinoids are both constitutively synthesized and induced by insect feeding, both *IGPS1* and *IGPS3* could function in W22 benzoxazinoid biosynthesis. In contrast to *IGPS1* and *IGPS3*, *IGPS2* expression was unchanged or even slightly decreased by insect feeding in both W22 and B73 (Figure 4; Figure S2; Tzin *et al*., 2015, Tzin *et al*., 2017).

**Figure 4.**
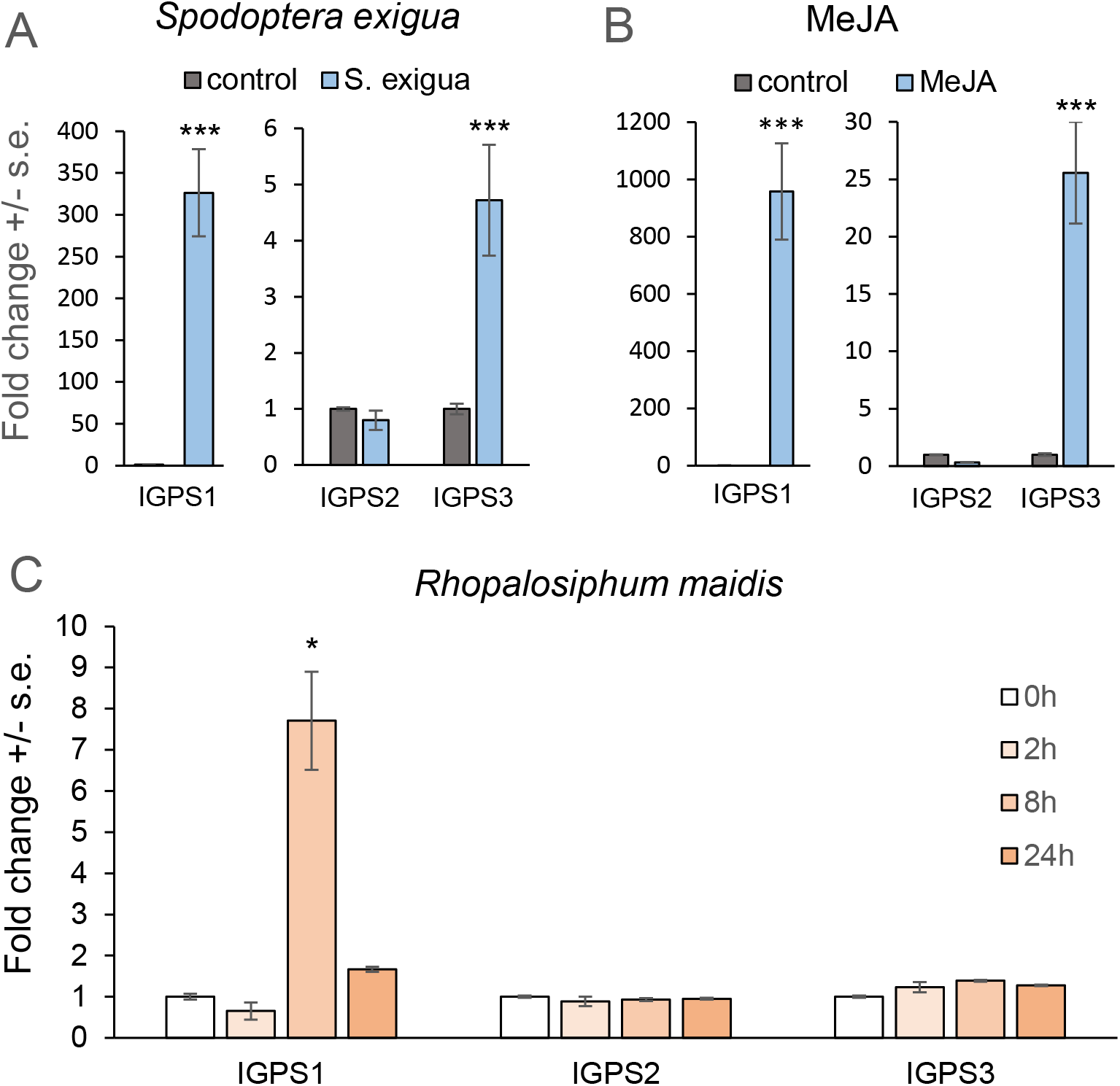
Transcript abundance of *IGPS1, IGPS2*, and *IGPS3* in maize inbred line W22 leaves. Transcript levels of maize *IGPS* genes were determined after (A) 24 h after initiation of *Spodoptera exigua* feeding (mean +/- s.e. of n = 5), (B) 24 h after methyl jasmonate induction (mean +/- s.e. of N = 3), and (C) *Rhopalosiphum maidis* infestation (mean +/- s.e. of N = 8). Values are fold change relative to uninfested controls (n = 5). *P < 0.05, ***P < 0.001 by Student’s *t*-test (panels A and B) or Dunnett’s test relative to 0 h control (panel C).

### Transposon insertions in *IGPS1* and *IGPS3* reduce benzoxazinoid levels

A search of publicly available transposon insertions (McCarty *et al*., 2005, Settles *et al*., 2007, Williams-Carrier *et al*., 2010) identified mu-02540, a likely *IGPS1* knockout line with a *Mu* insertion in the second exon (Chr.7: 82883253 bp to 82883261 bp; maize W22 genome v2). We used quantitative reverse transcriptase-PCR (QRT-PCR) to compare homozygous, heterozygous, and wildtype plants after methyl jasmonate induction (Figure 5A). Heterozygous plants showed lower expression levels than wildtype, and no *IGPS1* transcripts above background levels could be amplified from the homozygous mutant plants. The expression of *IGPS3*, but not *IGPS2*, was significantly increased in the *igps1* mutant line (Figure 5B).

**Figure 5.**
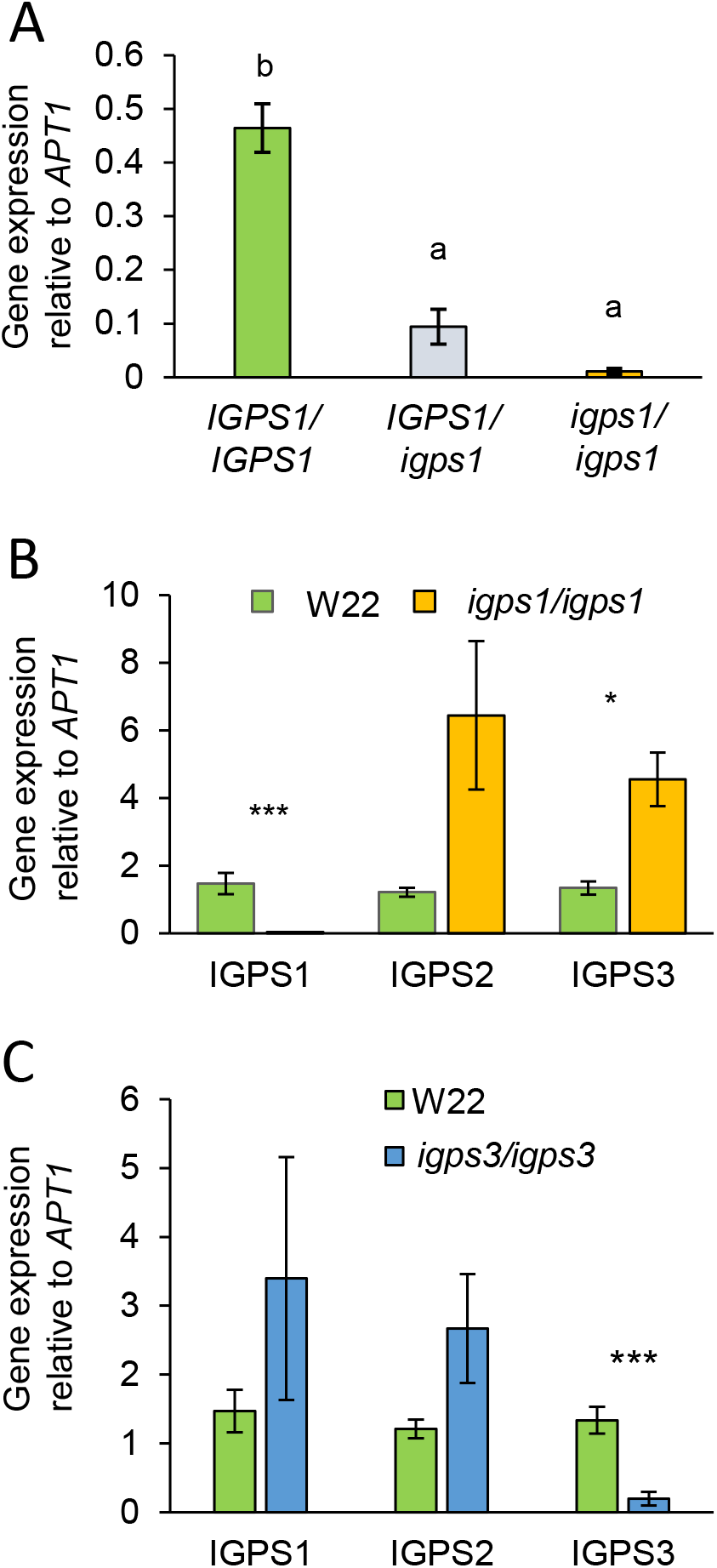
Effects of *igps1* and *igps3* knock-out mutations on gene expression. (A) *IGPS1* expression in a segregating population after induction with methyl jasmonate (mean +/- s.e. of N = 6). (B) Expression of *IGPS* genes in wildtype W22 and a homozygous *igps1* mutant line after methyl jasmonate induction (mean +/- s.e of N = 6). (C) *IGPS* gene expression in wildtype W22 and a homozygous *igps3* mutant line after induction with methyl jasmonate (mean +/- s.e. of N = 4). Different letters indicate P < 0.05, ANOVA followed by Tukey’s HSD test. *P < 0.05, ***P < 0.001 by Student’s *t*-test.

We identified a likely *IGPS3* knockout mutation in the *Mu-*Illumina collection (mutant line 228894.6; chr10: 69445669 - 69446382; Williams-Carrier *et al*., 2010). Plants were induced with methyl jasmonate to activate *IGPS3* transcription, and qRT-PCR was used to verify homozygous mutants and wildtype plants. *IGPS3* expression was significantly reduced (p < 0.001) in the mutant line, but the other two *IGPS* genes showed no significant changes in expression levels (Figure 5C).

To investigate IGPS2 involvement in benzoxazinoid or indole formation, we characterized several maize lines with transposon insertions upstream of the *IGPS2* open reading frame: *Mu*-Illumina_235844.6 (chr.10: 124084078-124084606; v3), *Mu*_Illumina_252733.6 (chr.10: 124083942-124084621; v3), *Mu*_Illumina_226348.6 (chr.10: 124084282-124084696; v3); *Mu*_Illumina_243642.6 (chr.10: 124084273-124084590; v3), UfMu_mu1030978 (Chr.10: 124084219-124084227; v3) and UfMu_mu1013033 (Chr.10: 124084306-124084314; v3) (McCarty *et al*., 2005, Settles *et al*., 2007, Williams-Carrier *et al*., 2010). However, none of these transposon insertions reduced *IGPS2* transcript levels (data not shown). No *Mu* or *Ds* transposon insertions in the coding region of *IGPS2* were available in public collections.

Both heterozygous and homozygous *igps1* mutant plants had significantly lower levels of DIMBOA (2,4-dihydroxy-7-methoxy-1,4-benzoxazin-3-one), HMBOA (2-hydroxy-7-methoxy-2H-1,4-benzoxazin-3(4H)-one), DIMBOA-Glc (2-O-β-D-glucopyranosyloxy-4-hydroxy-7-(2H)-methoxy-1,4-benzoxazin-3(4H)-one), DIM2BOA-Glc (2-O-β-D-glucopyranosyloxy-4-hydroxy-7,8-dimethoxy-(2H)-1,4-benzoxazin-3(4H)-one) and HM2BOA-Glc (Glc-2-O-β-D-glucopyranosyloxy-7,8-dimethoxy-(2H)-1,4-benzoxazin-3(4H)-one) than wildtype plants after methyl jasmonate treatment (Figure 6A). The abundance of all measured benzoxazinoids was decreased in homozygous *igps3* mutant plants (Figure 6B), though less so than in the *igps1* mutant line. The level of methyl jasmonate-induced indole accumulation was lower in both *igps1* and *igps3* mutant lines (Figure 6C). By contrast, there were no significant changes in the tryptophan content in either of the mutant lines relative to wildtype W22 (p > 0.05, *t-*test; Figure 6D,E).

**Figure 6.**
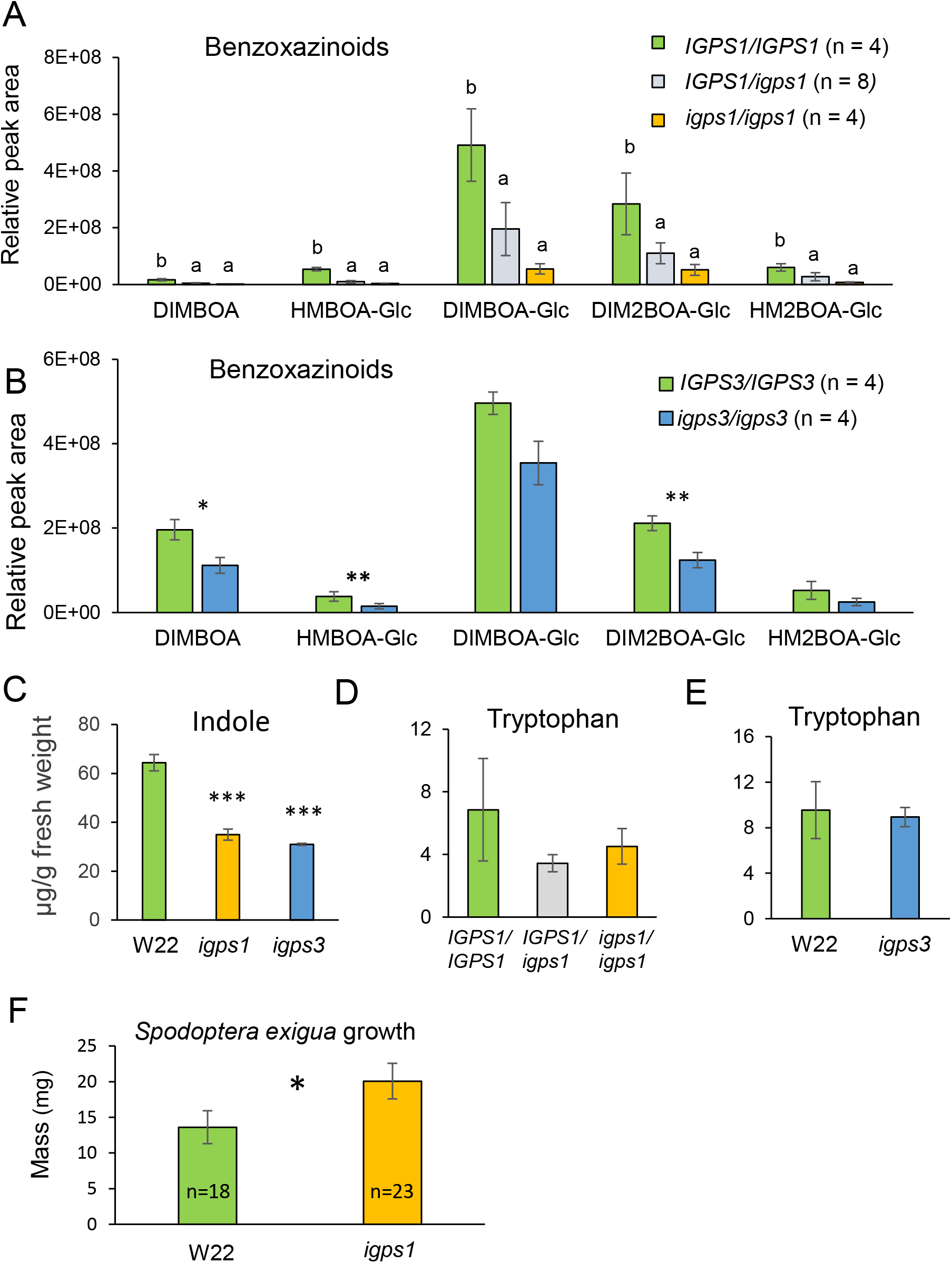
Effects of *igps1* and *igps3* knock-out mutations on metabolites and caterpillar growth. (A) Comparison of the five most abundant benzoxazinoids in an *igps1* segregating population (mean +/- s.e of N = 4-8). (B) The five most abundant benzoxazinoids in *igps3* mutant and wildtype plants (mean +/- s.e. of N = 4). (C) Tryptophan abundance in an *igps1* segregating population (mean +/- s.e. of N = 4-8). (D) Tryptophan in *igps3* mutant and wildtype plants (mean +/- s.e. of N = 4). (E) Free indole in wildtype, *igps1*, and *igps3* plants (mean +/- s.e. of N = 4). (F) Growth of *Spodoptera exigua* larvae on homozygous *igps1* mutant and wildtype W22. *P < 0.05, **P < 0.01, \*\*\**P* < 0.001 by Student’s *t*-test. Different letters on bars indicate P < 0.05, ANOVA and Fischer’s LSD test. No significant differences were detected for panels D and E (P > 0.05). Samples in panels A-E were treated methyl jasmonate to induce production of defensive metabolites.

*Spodoptera exigua* is a generalist lepidopteran herbivore that grows better on *bx1::Ds* and *bx2::Ds* mutant plants, which are almost completely devoid of benzoxazinoids, than on wildtype W22 maize (Tzin *et al*., 2017). We conducted bioassays to determine whether the benzoxazinoid decrease in the homozygous *igps1* mutant line (Figure 6A) also affects *S. exigua* caterpillar growth. There was a significant increase in caterpillar body mass, with an average weight of 20 mg on homozygous *igps1* mutant plants compared to 13.6 mg average weight on wildtype plants after nine days of feeding (p < 0.05, *t-*test; Figure 6F). We did not expect to see a measureable effect on caterpillar growth due to the relatively small decrease in benzoxazinoids in the *igps3* mutant (Figure 6B) and therefore did not conduct experiments with this line.

### IGPSs enzymes are localized in maize chloroplasts

In addition to gene expression levels, another factor that can influence the function of IGPS enzymes is their sub-cellular localization. The TargetP 1.1 software program (http://www.cbs.dtu.dk/services/TargetP/), which predicts subcellular localization and cleavage sites for signal peptides, indicated a chloroplast location for all three maize IGPS proteins. To experimentally confirm this, we cloned the *IGPS* open reading frames (B73-alleles) into the pEXSG vector (Bethke *et al*., 2009), to introduce an *C*-terminal yellow fluorescent protein (YFP) tag. These constructs were transiently expressed in maize protoplasts and analyzed by confocal microscopy. Positive transformed cells containing the *C*-terminal tagged protein showed a YFP signal after excitation with a 514 nm laser. Protoplast transformation of *pEXSG::IGPS1-YFP, pEXSG::IGPS2-YFP*, and *pEXSG::IGPS3-YFP* revealed similar YFP signals in maize chloroplasts (Figure 7A), with the YFP fluorescence overlapping the red chlorophyll autofluorescence of the plastids. Previous research with control constructs expressed in protoplasts showed that YFP by itself was located in the cytosol (Richter *et al*., 2016).

**Figure 7.**
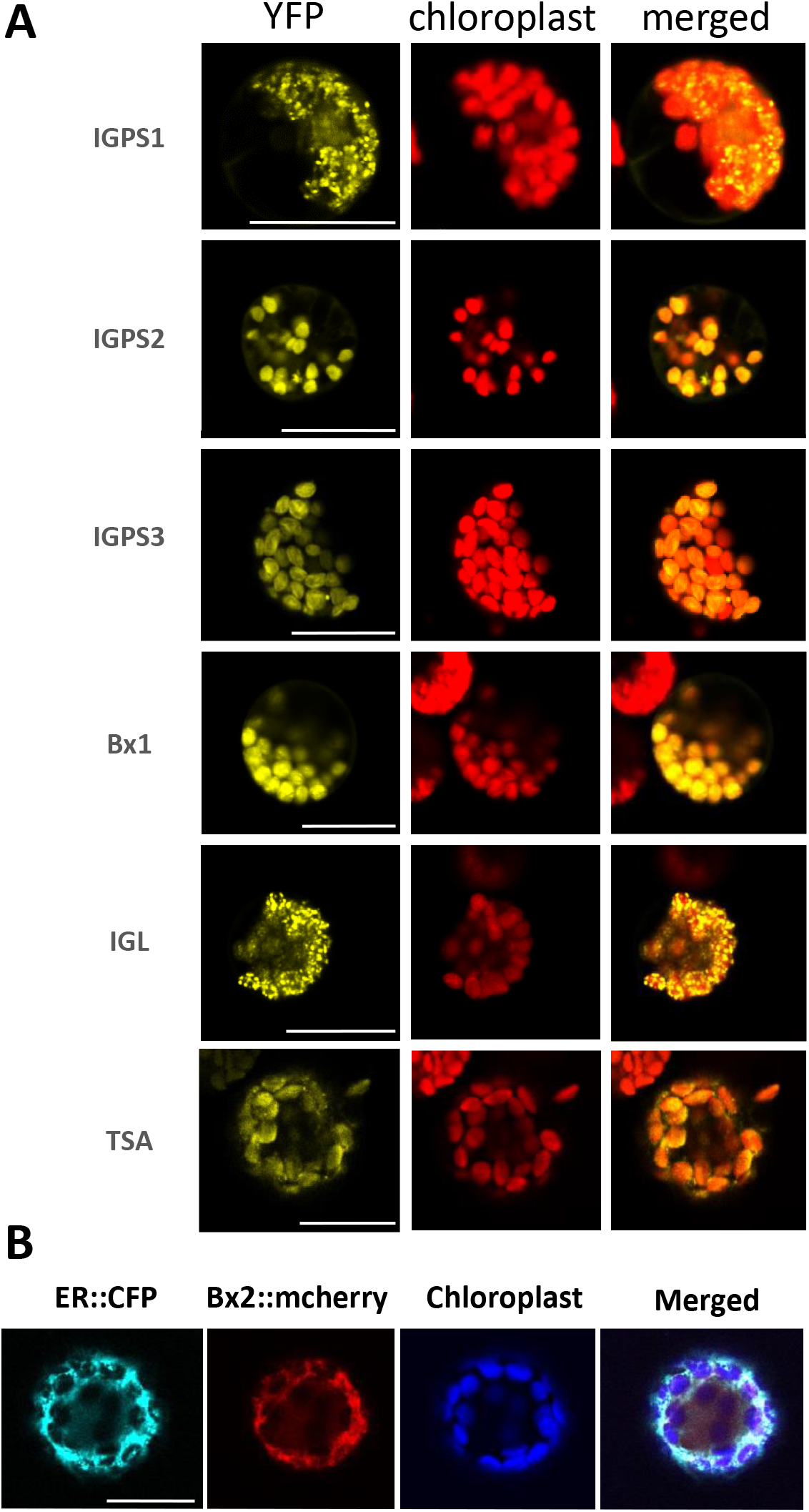
Subcellular localization of IGP synthases, BX1, BX2, TSA, and IGL. **(A)** Images of maize mesophyll protoplasts transiently expressing *35S-IGPS1-YFP, 35S-IGPS2- YFP, 35S-IGPS3-YFP, 35S-YFP-BX1, 35S-YFP-IGL*, and *35S-YFP-TSA*, showing YFP fluorescence in chloroplasts, chlorophyll auto-fluorescence, and merged images. **(B)** Colocalization of *35S-BX2-mcherry* with *ER-CFP*. The white scale bars indicate 25 μm.

In addition to the IGPS proteins, we determined the sub-cellular localization of IGL, TSA, BX1, and BX2 (Figure 1, Figure 7A). As has been reported previously for TSA-GFP (Kriechbaumer *et al*., 2008), the protein products of *pENSG::YFP-BX1, pENSG::YFP-IGL*, and *pENSG::YFP-TSA* were located to the plastids of transformed maize protoplasts (Figure 7A). By contrast, BX2 carrying an *N*-terminal mCherry tag (Karimi *et al*., 2016) was co-localized with an endoplasmic reticulum marker, rather than to the chloroplasts (Figure 7B).

### Interactions of IGPS and indole-3-glycerolphosphate lyases

To determine whether the three IGPS enzymes interact with BX1, TSA, and IGL, we performed BiFC interaction studies for all nine possible two-protein combinations. *IGPS1, IGPS2*, and *IGPS3* were cloned into p2YC (Yang *et al*., 2007), such that their C-termini were fused to a C-terminal YFP fragment (amino acids 158-238). The open reading frames for *BX1*, *TSA*, and *IGL* were cloned into p2YN (Yang *et al*., 2007), fusing an *N*-terminal YFP fragment (amino acids 1-159) at the *C*-termini. All constructs were transformed into *Agrobacterium tumefaciens* and infiltrated into *Nicotiana benthamiana* plants in pairs. Two days after infiltration, protoplasts from the infiltrated leaf area were isolated and analyzed by confocal microscopy. Strong YFP signals were localized in the chloroplasts for the combinations of IGPS1-C/Bx1-N and IGPS1-C/IGL-N (Figure 8A). On the other hand, coexpression of IGPS1-C and TSA-N did not result in reconstitution of YFP fluorescence. By contrast, coexpression of IGPS2 with TSA resulted in a strong fluorescence signal, and weaker signals were observed when IGPS2 was coexpressed with BX1 or IGL (Figure 8B). IGPS3 consistently showed reconstitution of the fluorescence when coexpressed with all three indole-3-glycerolphosphate lyase partners (BX1, IGL, and TSA; Figure 8C). Additionally, by cloning *IGPS2* and *IGPS3* cDNA sequences into p2YN, we were able to test whether IGPS1 could interact with the other two IGPS enzymes. The co-expression of IGPS1 and IGPS3 could be detected, suggesting a possible complex that could bind to the BX1 and IGL proteins (Figure 8D). However, no interaction signal was detected for the IGPS1 - IGPS2 combination.

**Figure 8.**
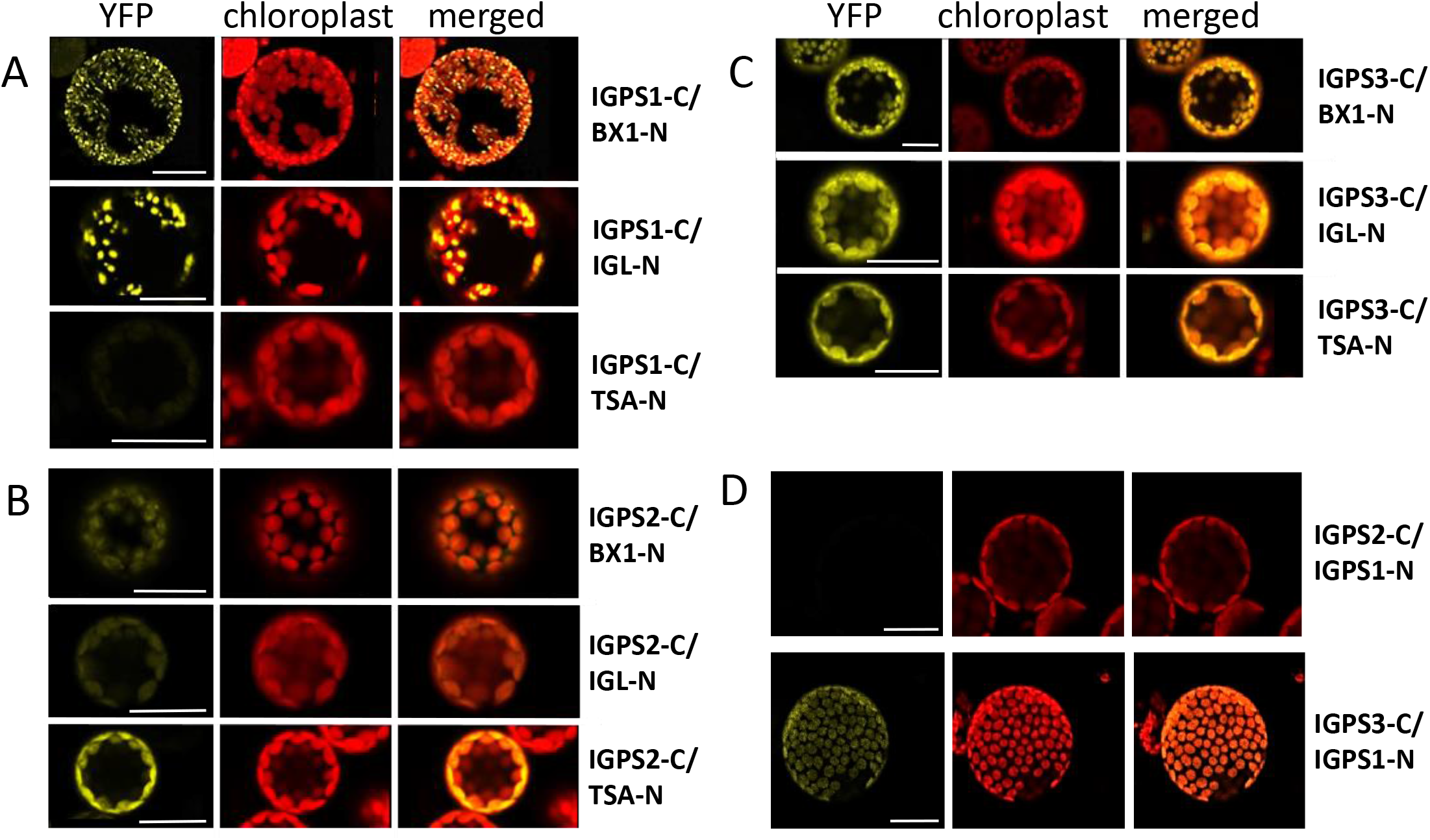
Interactions of IGPS proteins with BX1, TSA, and IGL by BiFC assays in *N. benthamiana*. The overlap of YFP (yellow) and the chloroplast auto-fluorescence (red) images is shown in the column labeled merged. IGPSs were fused with C-terminus of YFP (c-YFP) and each construct was co-transformed into *N. benthamiana* leaves, with BX1, TSA, and IGL, which were fused with the N-terminus of YFP (n-YFP). (A) Interactions of IGPS1 with BX1, TSA, and IGL, (B) interactions of IGPS2 with BX1, TSA, and IGL, (C) interactions of IGPS2 with BX1, TSA, and IGL, and (D) interactions of IGPS1 N-terminal YFP fusions with IGPS2 and IGPS3 C-terminal YFP fusions. The white scale bars indicate 25 μm.

## Discussion

Both complementation of an *E. coli trpC* mutation and analysis of maize transposon insertion lines indicated that maize *IGPS1, IGPS2*, and *IGPS3* encode functional indole-3-glycerolphosphate synthases. In a phylogenetic analysis, IGPS2 clustered with previously characterized Arabidopsis IGPS proteins (Li *et al*., 1995a; Figure 2), suggesting a role in primary metabolism. By contrast, IGPS1 and IGPS3 are in separate phylogenetic clades, consistent with their functions in defensive benzoxazinoid metabolism that is not found in Arabidopsis and other dicots that are included in the dendrogram. It is possible that the IGPS1 and IGPS3 enzyme clades in other monocots (rice, wheat, sorghum, barley and *S. viridis*; Figure 2) have additional functions in the production of defense-related metabolites in these species. A similar phylogenetic analysis of maize indole-synthesizing enzymes showed that BX1 and IGL, which contribute to defensive metabolism, cluster separately from the Arabidopsis and maize TSA proteins (Frey *et al*., 2000, Dutartre *et al*., 2012). Together, these observations are consistent with the hypothesis that pathways for the production of defense-related benzoxazinoids and indole arose through the duplication and repurposing of ancestral IGPS and TSA enzymes that contribute to primary metabolism in maize and other plants.

Our results are consistent with the observations of Wisecaver *et al*. (2017), who used global co-expression network analysis to propose that *IGPS1* is part of the benzoxazinoid biosynthesis pathway in maize. Both B73 and W22 inbred lines show a greater than a 100-fold increase of *IGPS1* transcripts in response to caterpillar feeding (Figure 4A, Figure S2A, Tzin *et al*., 2017). Consistent with this induction, *igps1* maize mutants have lower levels of benzoxazinoids (Figure 6A), and *S. exigua* caterpillars grow better on *igps1* mutant than on wildtype W22 plants (Figure 6F). *IGPS3* did now show up in co-expression studies with *BX* genes (Wisecaver *et al*., 2017), perhaps due to the lower level of defense-related induction of gene expression, which may have been a factor in the integrative comparison of multiple gene expression studies.

Unlike in the case of IGPS1, IGPS2, IGPS3, BX1, IGL, and TSA, which localize to the chloroplasts when overexpressed in maize protoplasts, we were not able to verify a chloroplastic location for BX2, a cytochrome P450 that catalyzes the first oxygenation reaction in the pathway from indole to DIBOA-Glc (Frey *et al*., 1995). Consistent with localization of the BX2 enzymatic activity in microsomes after cell separation (Glawischnig *et al*., 1999), our protein fusion studies indicated that BX2 is localized in the endoplasmic reticulum (Figure 7B). Given that both *bx1* and *bx2* mutations result in an almost complete loss of benzoxazinoids in maize inbred line W22 (Betsiashvili *et al*., 2015, Tzin *et al*., 2015), there must be a mechanism by which the indole that is produced by BX1 in the chloroplast is passed to BX2 in the endoplasmic reticulum. Indole produced by TSA or IGL appears to be in a separate metabolic pool and is not used as a substrate by BX2. There are direct interactions between maize TSA and TSB enzymes (Kriechbaumer *et al*., 2008), which may prevent the release of free indole in this pathway. However, as BX1 and BX2 are in different cellular compartments (chloroplast and endoplasmic reticulum, respectively), it is harder to imagine a direct interaction that allows passage of indole from one protein to the other, without including free indole that is produced by IGL.

Indole production highly induced in maize after herbivore feeding (Frey *et al*., 2000, Erb *et al*., 2015). As in the case of benzoxazinoids, free indole was less abundant in both *igps1* and *igps3* mutant lines. Whereas *bx1* mutations cause an almost complete loss of benzoxazinoids (Tzin *et al*., 2017), and IGL is primarily involved in indole biosynthesis (Frey *et al*., 2000), IGPS1 and IGPS3 seem to be involved in both metabolic pathways. Consistent with a function in both benzoxazinoid and free indole biosynthesis, IGPS1 and IGPS3 appear to interact with both BX1 and IGL (Figure 8A,C), and interactions between IGPS1 and IGPS3 (Figure 8D) may also contribute to the overlapping functions of these enzymes. An *igps1 igps3* double mutant would be required to determine whether benzoxazinoids and free indole in maize are derived from these two enzymes alone or whether IGPS2 also functions in the production of defensive maize metabolites.

Among the three maize indole-3-glycerolphosphate lyases, IGPS2 had the strongest interaction with TSA (Figure 8B), which is thought to function in tryptophan biosynthesis (Figure 1). Consistent with the hypothesis that tryptophan is synthesized primarily by an IGPS2-dependent pathway, neither *igps1* nor *igps3* mutations significantly decreased the tryptophan content of maize leaves (Figure 6D,E). Maize *tsb1 tsb2* double mutations caused a seedling lethal phenotype (Wright *et al*., 1992), and it has been hypothesized that TSA, the only maize indole-3-glycerolphosphate lyase that interacts with TSB1, is therefore also essential (Kriechbaumer *et al*., 2008). By extension, if IGPS2 has a primary function in providing IGP to TSA, it may also be essential. When conducting our experiments, we were not able to find any *IGPS2* transposon knockout mutations of *IGPS2* in public collections. However, there are predicted *IGPS2* coding-region insertions in the recently published BonnMu collection (Marcon *et al*., 2020). If these insertions can be confirmed, it will be possible to determine whether IGPS2 has an essential function in maize metabolism.

BX1, IGL, and TSA channel their metabolic products to particular pathways. The free indole-producing function of maize IGL (Frey *et al*., 2000, Ahmad *et al*., 2011) is not provided by the BX1 or TSA enzyme activity. Similarly, a *bx1* knockout phenotype cannot be complemented by the activity of IGL or TSA, only by supplementing free indole (Frey *et al*., 1997, Melanson *et al*., 1997). To this metabolic channeling, we can now add the effects of IGPS1, IGPS2, and IGPS3, which have distinct, though partially overlapping, functions in channeling IGP to specific indole pools that are formed by BX1, IGL, and TSA. The protein interactions that we have observed (Figure 8) provide new insight into the regulation of indole metabolism and how, within the chloroplast, separate pools of this metabolite can be used for primary metabolism (tryptophan) and specialized metabolism (volatile indole and benzoxazinoids).

## Materials & methods

### Plant material and methyl jasmonate treatment

Maize (*Zea mays* L.) seeds from the inbred lines B73, W22, and transposon insertion lines Ufmu-02540, UfMu_mu1030978, UiMumul0l3033, UfMu_mu1030978 (http://maizecoop.cropsci.uiuc.edu/) and *Mu*-Illumina_228854.6, *Mu*-Illumina_235844.6, *Mu-* Illumina_252733.6, *Mu*-Illumina_226348.6, *Mu*-Illumina_235844.6, *Mu*-Illumina_252733.6, *Mu*-Illumina_226348.6, *Mu*-Illumina_243642.6, UfMu_mu1030978 (Barkan lab, http://molbio.uoregon.edu/barkan/) were germinated in water-filled Petri dishes for three days and were then transferred into plastic pots filled with a soil mix (0.16 m3 Metro-Mix 360 from Scotts, Marysville, OH, USA; 0.45 kg finely ground lime; 0.45 kg Peters Unimix from Griffin Greenhouse Supplies, Auburn, NY, USA; 68 kg Turface MVP from Banfield-Baker Corp., Horseheads, NY, USA; 23 kg coarse quartz sand, and 0.018 m3 pasteurized field soil). Plants were grown for about 2 weeks in climate-controlled chambers (16 h light:8 h dark cycle, 180 mmol photons m^-2^ s^-1^ light intensity at constant 23 °C and 60% humidity). For methyl jasmonate induction experiments using a previously described method (Richter *et al*., 2015, Tamiru *et al*., 2017), the third leaf of each plant was cut off and incubated in 2 ml tap water with 250 μM methyl jasmonate (Sigma-Aldrich; 392707) for 24 h, after which the leaves were snap-frozen in liquid nitrogen and stored at −80 °C until they were used for assays. Control leaves were harvested directly into liquid nitrogen without methyl jasmonate treatment.

### Gene expression induction by insect feeding

Maize inbred line W22 gene expression assays after *R. maidis* infestation were performed as described for inbred line B73 by Tzin et al. (2015), but we used 10-day-old seedlings and samples were harvested at different time points: 0, 2, 8, and 24 h. The control plants received empty cages without aphids for 24 h and were harvested and frozen in liquid nitrogen at the same time as the experimental samples. To induce gene expression changes in maize line W22 with *S. exigua* caterpillars, we followed the protocol of Tzin *et al*. (2017). Leaf tissue for analysis was harvested after 24 h of caterpillar feeding.

### Total RNA extraction and gene expression analysis

Harvested leaf material was frozen in liquid nitrogen, and ground to a fine powder for RNA isolation with on-column DNA digestion, using the SV Total RNA isolation kit (Promega, Madison, WI, USA). The total RNA concentration was measured with a NanoDrop instrument (2000c; ThermoFisher Scientific Inc., Waltham, MA, USA), and 500 ng total RNA was incorporated for first-strand cDNA synthesis using the M-MLV reverse transcriptase kit (TaKaRa Bio USA, Mountain View, CA, USA), and the library was used as a template for qRT-PCR analysis. Thereafter, 5 μl SYBR Green Mix (Universal SYBR Green Supermix, Bio-Rad Laboratories, Hercules, CA), 1 μl gene-specific forward primer, 1 μl gene-specific reverse primer, 0.5 μl template, and 2.5 μl PCR-grade water were mixed. The PCR reaction was performed in a QuantStudio™ 6 Flex Real-Time PCR system instrument (384-well, Applied Biosystems, Foster City, CA) with an initial incubation at 95°C for 10 min. The following cycle was repeated 40 times: 95°C for 30 s, 60°C for 15 s, and 72°C for 30 s. Maize adenine phosphate transferase 1 (*APT1*) was used as a reference gene. For each analyzed gene, a cDNA pool from all plants was diluted from 1 times to 80 times to generate a standard curve. With Applied Biosystems software, the ΔC_T_ (△-threshold cycle) for each gene was calculated relative to the reference gene. For the CT values an average from at least three biological replicates was calculated. The primer sets used to amplify the genes are given in Supplementary Table S1.

### Transcriptome sequencing and RNAseq data analysis

The transcriptome of the W22 aphid time course experiment was sequenced using the 3’RNAseq method (Kremling *et al*., 2018). RNA was isolated as described above with a total of 8 biological replicates for each treatment. The purity of all RNA samples was proofed with a NanoDrop2000 instrument (Thermo Scientific). The 3’RNA-seq libraries were prepared from 6 μg total RNA at the Cornell Genomics facility (http://www.biotech.cornell.edu/brc/genomics-facility; Kremling et al. 2018). Trimmomatic version 0.36 (Bolger *et al*., 2014) was used to remove the first 12 bp from each read. STARaligner v.2.4.2 (Dobin *et al*., 2013) was used to align the reads against the W22 genome (Springer *et al*., 2018), with a maximum of one location to map to (−outFilterMultimapNmax 1). Another step in the pre-processing was increasing the region size of each gene by 500 bp to decrease the number of genes mapped to a region with no feature from > 120000 to < 60000 on average across the libraries. For this, we used the slop command from bedtools (https://bedtools.readthedocs.io/en/latest/) with −r 500 (−r the number of base pairs to add to the end coordinate). The program Tablet (https://ics.hutton.ac.uk/tablet/) was used to visualize the annotated genes on each chromosome. HTSeq32 v.0.6.1 were used to obtain genelevel counts from the resulting BAM files (htseq-count −r name −s yes −a 10 −f bam). To normalize the read counts, the CPM (count per million) was calculated by dividing the read counts for each gene by the total counts of the RNAseq sample. Illumina sequencing data were submitted to the Sequence Read Archive (https://www.ncbi.nlm.nih.gov/sra) as data set SUB8291142 (Submission ID), PRJNA669140 (Bioproject ID).

### Isolation and cloning of cDNA

To isolate the open reading frames of *IGPS1, IGPS2, IGPS3, TSA, IGL, BX1*, and *BX2* from B73 cDNA, specific primers (Supplementary Table S1) were created based on the B73_v4 maize genome. iproof high fidelity polymerase (Bio-Rad) was used for PCR amplification. The fragments were cloned into the specific vectors mentioned for each experiment and sequenced for their accuracy.

### Complementation assays in *E. coli*

The open reading frames of *IGPS1, IGPS2* and *IGPS3*, including their respective signal sequences, were cloned into the bacterial expression vector pASKIBA37+ (IBAGmbH, Göttingen, Germany). To amplify these genes, special primers with a *BsaI* restriction site were created with the ‘‘Primer D’ Signer’’ (IBA GmbH) (Supplementary Table S1). After sequencing the constructs, plasmids were used for complementation assays in the *E. coli trpC9800*^-^ tryptophan-auxotrophic mutant background. The plasmids were transformed into *trpC9800*^-^ competent cells and plated out on 100 μg/ml ampicillin LB-agar plates. To test their activity in the mutant background, a starter overnight culture of 2 ml LB medium with 100 μg/ml ampicillin was prepared for each construct, including the empty vector as a negative control. The next morning, the starter culture was used to inoculate 5 ml of tryptophan-deficient Vogel and Bronner (Vogel and Bonner, 1956) minimal medium containing 2 % dextrose and 100 μg/ml ampicillin. For positive controls, 20 μg/ml *L*-tryptophan was added. All cultures started with an optical density at 600 nm (OD600) = 0.2 in the time-course experiment. OD600 was measured six times over 11 hours, until the bacteria had reached stationary phase.

### Indol-3-glycerolphosphate lyase phylogenetic tree

A dendrogram analysis was conducted using MegaX software (Kumar *et al*., 2018). For the alignment we used the MUSCLE Codon algorithm (gap open −2.9, gap extend 0, hydrophobicity multiplier 1.2, clustering method, upgma). The evolutionary history was inferred by using the Maximum Likelihood method and JTT matrix-based model (Jones *et al*., 1992). The percentage of trees in which the associated taxa clustered together is shown next to the branches. Initial tree(s) for the heuristic search were obtained automatically by applying Neighbor-Join and BioNJ algorithms to a matrix of pairwise distances estimated using a JTT model, and then selecting the topology with superior log likelihood value. A discrete Gamma distribution was used to model evolutionary rate differences among sites (5 categories (+*G*, parameter = 0.7732)). In an analysis involved 27 amino acid sequences, a bootstrap resampling analysis with 1,000 replicates was performed to evaluate the tree topology. All positions with less than 80% site coverage were eliminated, i.e., fewer than 20% alignment gaps, missing data, and ambiguous bases were allowed at any position (partial deletion option). There were a total of 358 positions in the final dataset. The maize TSA protein was used as outgroup for the phylogeny.

### Metabolite analysis *via* LC-MS and data processing

Frozen, powdered leaf material was weighed (~100 mg), three volumes of 100% methanol were added to each sample, and samples were incubated for 45 min shaking at 4 °C. After a 10 min centrifugation step (15,000 x g) the samples were filtered in a 96-well MultiScreen filter plate (0.65 μm, Millipore, Burlington, MA). Reverse-phase liquid chromatography was performed using a DionexUltimate 3000 Series LC system (HPG-3400 RS High-Pressure pump, TCC-3000RScolumn compartment, WPS-3000TRS autosampler) controlled by ChromeleonSoftware (Thermo Fisher Scientific, Waltham, MA) and coupled to Orbitrap Q-Exactive mass spectrometer controlled by the Xcalibur software (Thermo Fisher Scientific). A Titan™ C18 UHPLC Column, 1.9 μm 100 mm × 2.1 mm at 40C and the flow rate of 0.5 ml/min of mobile phases A (H2O:0.1% formic acid) and B (AcN:0.1% formic acid) was used for the separation of target molecular features. The gradient starting condition was 0% B at 0 min, rising to 95% B in 15 min, then was held for 0.5 min, and followed by 0.5 min re-equilibration to the starting condition. A heated electrospray ionization source (HESI-II) in negative mode was used for the ionization, with the parameters of spray voltage at 3.5 kV, the capillary temperature at 380 C, sheath gas and auxiliary gas flow at 60 and 20 arbitrary units, respectively, S-Lens RF Level 50, and probe heater temperature 400°C. The data were acquired in the m/z range of 100-900, 140,000 FWHM resolution (at m/z 200), AGC target 3e6, maximum injection time of 200ms, in profile mode. Metabolite quantification was estimated with ThermoScientific Xcalibur™ Version 4.1.31.9 (Quan/Processing). For data analysis, the raw mass spectrometry files were converted to mzxml formats using the MSConvert tool (Chambers *et al*., 2012). The raw mzxml files were further processed using the XCMS (http://metlin.scripps.edu/download/, Tautenhahn *et al*., 2012) and the CAMERA software packages for R (http://www.bioconductor.org/packages/release/bioc/html/CAMERA.html). Data were analyzed using the XCMS-CAMERA mass scan data processing pipeline (Tautenhahn *et al*., 2008, Benton *et al*., 2010, Kuhl *et al*., 2012).

To quantify tryptophan content of maize leaf samples, a stock solution was made by diluting 1 M solution of *L-*tryptophan (Sigma-Aldrich, St. Louis, MO) in water 1:10 with methanol. Further dilutions were made using 100% methanol and a standard curve was prepared by LC-MS analysis, as described above, of samples containing to 0.3 to 10 ng/μl tryptophan (Supplemental Figure S3A). The concentration of tryptophan in maize tissue samples (μg/g) was calculated based on comparisons to this standard curve.

### Indole quantification by GC-MS

For the quantification of indole, leaf material was harvested, ground with a mortar and pestle in liquid nitrogen, and stored at −80°C until sample preparation. The volatile indole released from 50 mg powdered plant material was collected on a solid-phase microextraction (SPME) fiber (100 μm polydimethylsiloxane, fused silica 24 Ga; Supelco, Bellefonte, PA) for 45 min. Samples were analyzed using an Agilent 7890B gas chromatograph coupled to an Agilent 7000D triple quadrupole mass spectrometer (Agilent Technologies, Santa Clara, CA). Collection of volatiles from frozen and macerated tissue in this manner produces similar results to volatile collection from intact maize leaves (Köllner *et al*., 2004). To release volatiles from the fiber, an injection temperature of 220°C was used. Helium served as a carrier gas with a flow rate of 2.3 ml min^-1^ and nitrogen as collision gas with a flow rate of 1.5 ml min^-1^. To separate the volatiles, an EC5-MS column (30 m length, 0.25 mm diameter., and 0.25-μm film; Agilent J&W GC columns) was used under the following conditions: 60°C for 3 min, first ramp of 10°C min^-1^ to 150°C, second ramp of 100°C min^-1^ to 300°C, and final 2-min hold. Indole was identified and quantified by running purified indole (Supelco) as a standard. Indole was dissolved in dimethylsulfoxide (DMSO), diluted in water 1:10 and added to 50 mg powdered leaf tissue from uninduced maize inbred line W22, which contains no detectable indole. A standard curve of indole concentrations ranging from 0.58 μg to 5.8 μg indole added to 50 mg powdered leaf material (Supplemental Figure S3B), was used to quantify indole abundance in methyl jasmonate-induced leaf samples.

### Subcellular localization

Maize mesophyll protoplasts were isolated as described by Richter *et al*. (2016), with few modifications. After washing the protoplasts, the cells were transferred into the wash buffer instead of the MMG buffer, with a final concentration of ~ 1-2 x 10^6^/ ml.

The open reading frames of *IGPS1, IGPS2, IGPS3, BX1, BX2, TSA* and *IGL* were amplified without stop codons and cloned into pDONR 201 by BP clonase recombination (Invitrogen, Carlsbad, CA), before being transferred with a second recombination reaction (LR, Invitrogen) into the vector YFP-pENSg (Dissmeyer et al, 2017). To isolate high concentrations of each plasmid a Midi preparation (Qiagen, Hilden, Germany) was performed. All of the fusion proteins were transformed into maize mesophyll protoplasts as described above. After incubating maize protoplasts overnight, they were analyzed using the Leica TCS SP5 spectral imaging system (Leica Microsystems, Wetzler, Germany). The YFP fluorescence was excited with a 514 nm laser and the emission spectra were collected with a hybrid detector (HyD) at 523 nm to 591 nm. The autofluorescence of chloroplasts was excited with an argon laser and detected at 675 to 736 nm. Protoplasts from each transformation were imaged with a 20x water immersion objective. For localization of BX2 in the endoplasmic reticulum (ER), the open reading frame was cloned from pDONR 207 into pK7m34GW by LR Gateway recombination. For this reaction, the destination vector, pDONR, pENTR4-1 containing the 35S double promotor and plasmid pENTR2-3 with an mCherry tag were fused. Finally, the construct 35S:mCherry:BX2 was transformed into *A. tumefaciens* (GV3101) and grown overnight in LB medium containing appropriate antibiotics. Likewise, an ER-CFP (cyan fluorescent protein) marker construct (plasmid #953, Nelson *et al*., 2007) was transformed into *A. tumefaciens*. After mixing *A. tumefaciens* cultures of both constructs with the same OD600, they were co-infiltrated into *N. benthamiana* leaves. Two days after infiltration, protoplasts were isolated from the infiltrated leaf tissue area and used for microscopic analysis. CFP fluorophore excitation was performed using a 458 nm argon laser and detected at 468 to 496 nm. For co-localization, CFP and mCherry were scanned sequentially.

For interaction studies, the open reading frames of *IGPS1, IGPS2, IGPS3, TSA, IGL*, and *BX1* were cloned without stop codons into the BIFC vectors p2YC and p2YN (Kong *et al*., 2014). In addition, *IGPS1* was cloned into p2YN to study the interaction with IGPS2 and IGPS3. The constructs used for protein-protein interactions studies were co-infiltrated into *N. benthamiana* leaves, as described previously (DeBlasio *et al*., 2018). To detect the YFP signal and chloroplast autofluorescence, we used the microscope settings described above.

### Caterpillar growth assays

Eleven-day-old homozygous *igps1* mutant and wildtype W22 plants were used for caterpillar feeding assays. *Spodoptera exigua* eggs were purchased from Benzon Research (Carlisle, PA) and kept for 52 h in a 28°C incubator on artificial diet (Beet Armyworm Diet, Southland Products Inc., Lake Village, AR). One 10 x 15 cm organza mesh bag (amazon.com, item B073J4RS9C) containing one second instar *S. exigua* caterpillar was placed on each 11-day-old maize plant. The bags were gently closed with bobby pins without wounding the plant. After one week of feeding on maize plants, the caterpillars were weighed.

## Acknowledgements

This research was funded by a Deutsche Forschungsgemeinschaft fellowship to AR and Binational Agricultural Research and Development Agency (BARD) award US-4846-15C and US National Science Foundation award #2019516 to GJ. We thank Monika Frey for helpful advice and Kevin Ahern for assistance in identifying maize *IGPS* genes.

---

**Supplemental Figure S1.**
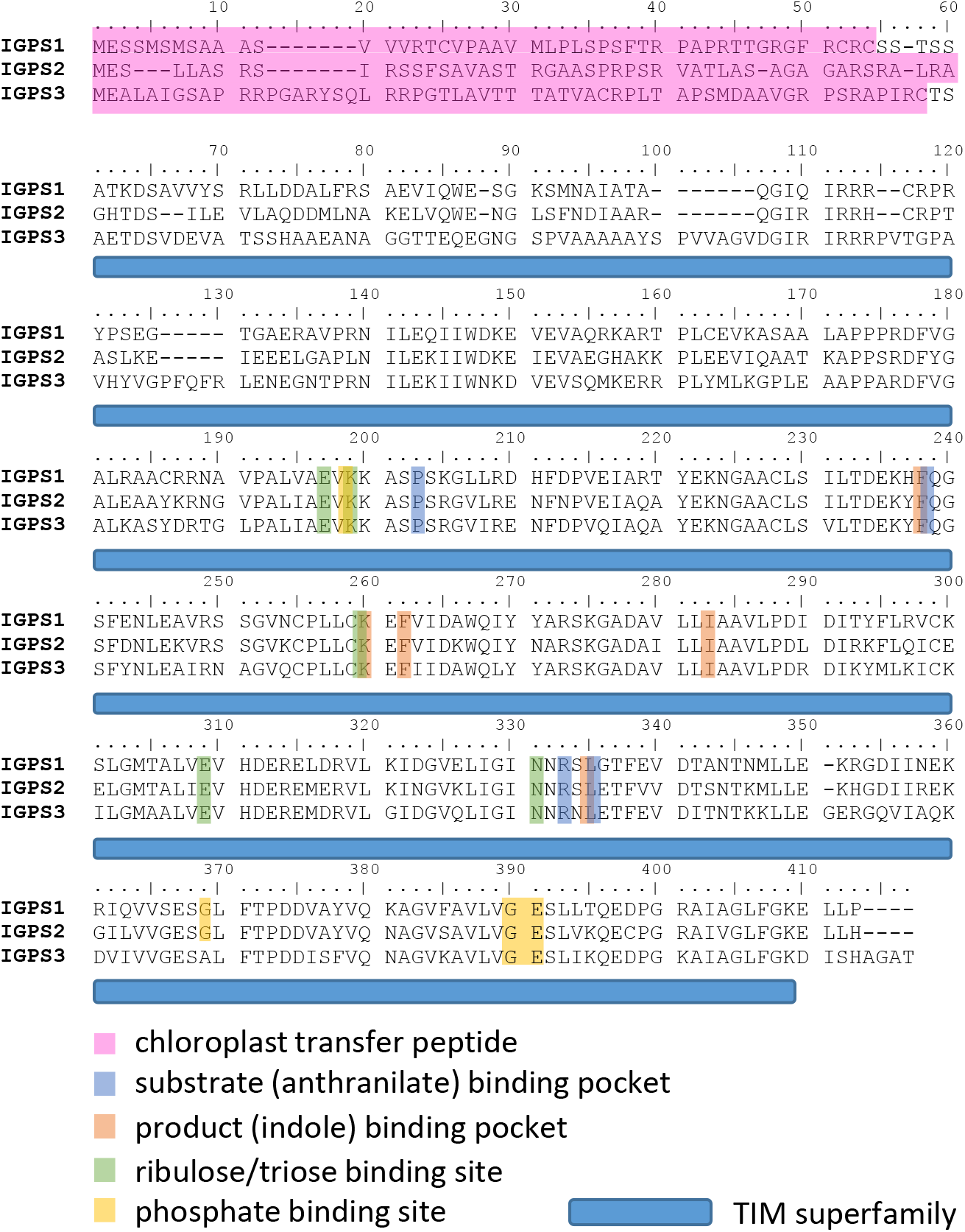
Alignment of the three maize IGPS proteins. Relevant protein features are highlighted.

**Supplemental Figure S2.**
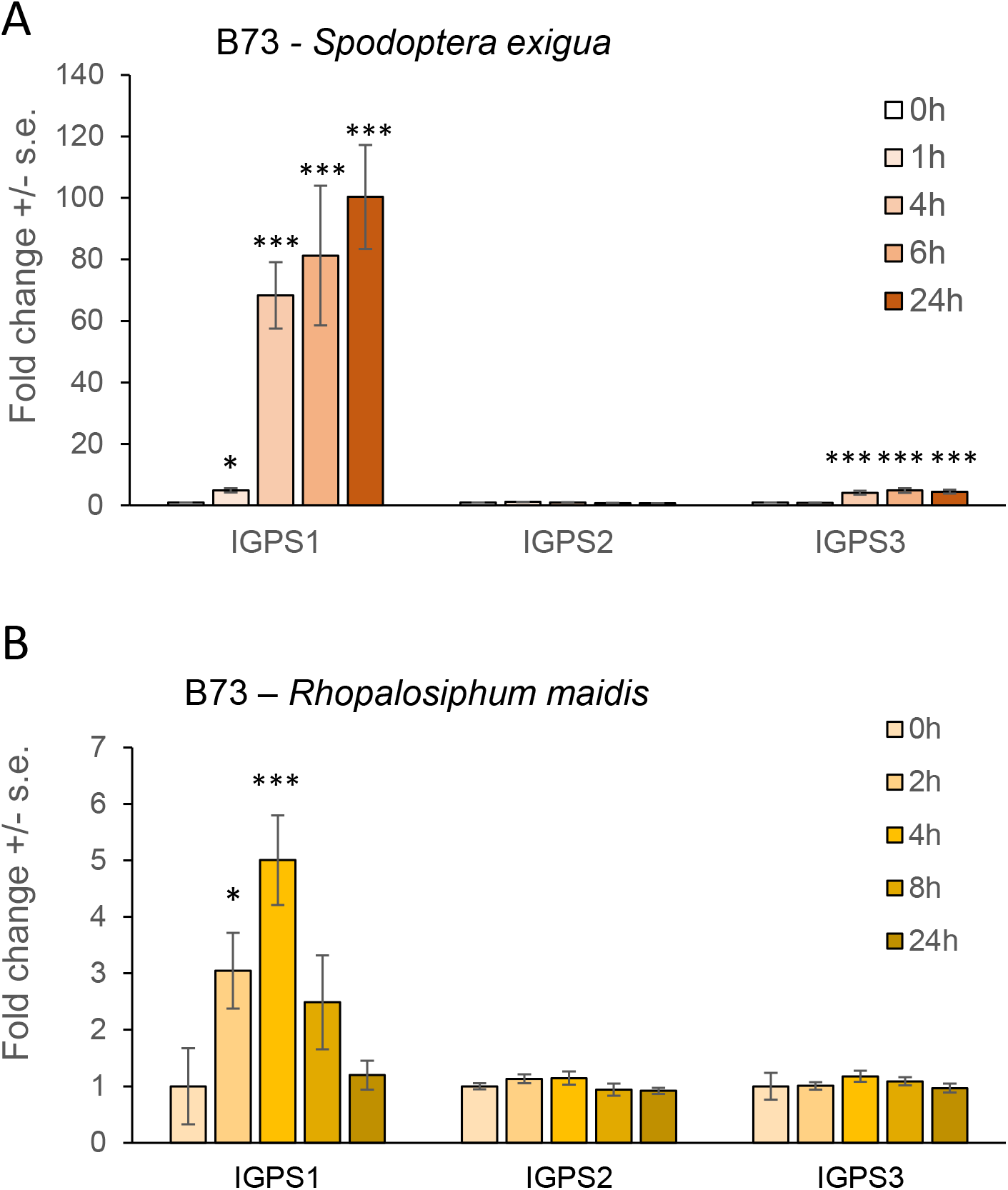
Expression of IGPS genes in response to insect feeding on maize inbred line B73. (A) Feeding by *Spodoptera* exigua (beet armyworm), mean +/- s.e. of N=4 (B) Feeding by *Rhopalosiphum maidis*, mean +/- s.e. of N = 5. *P < 0.05, **P < 0.01, ***P < 0.001, Dunnett’s test relative to 0-hr control. These figures are produced from RNAseq data presented in Tzin et al (2017) and Tzin et al (2015), respectively.

**Supplemental Figure S3.**
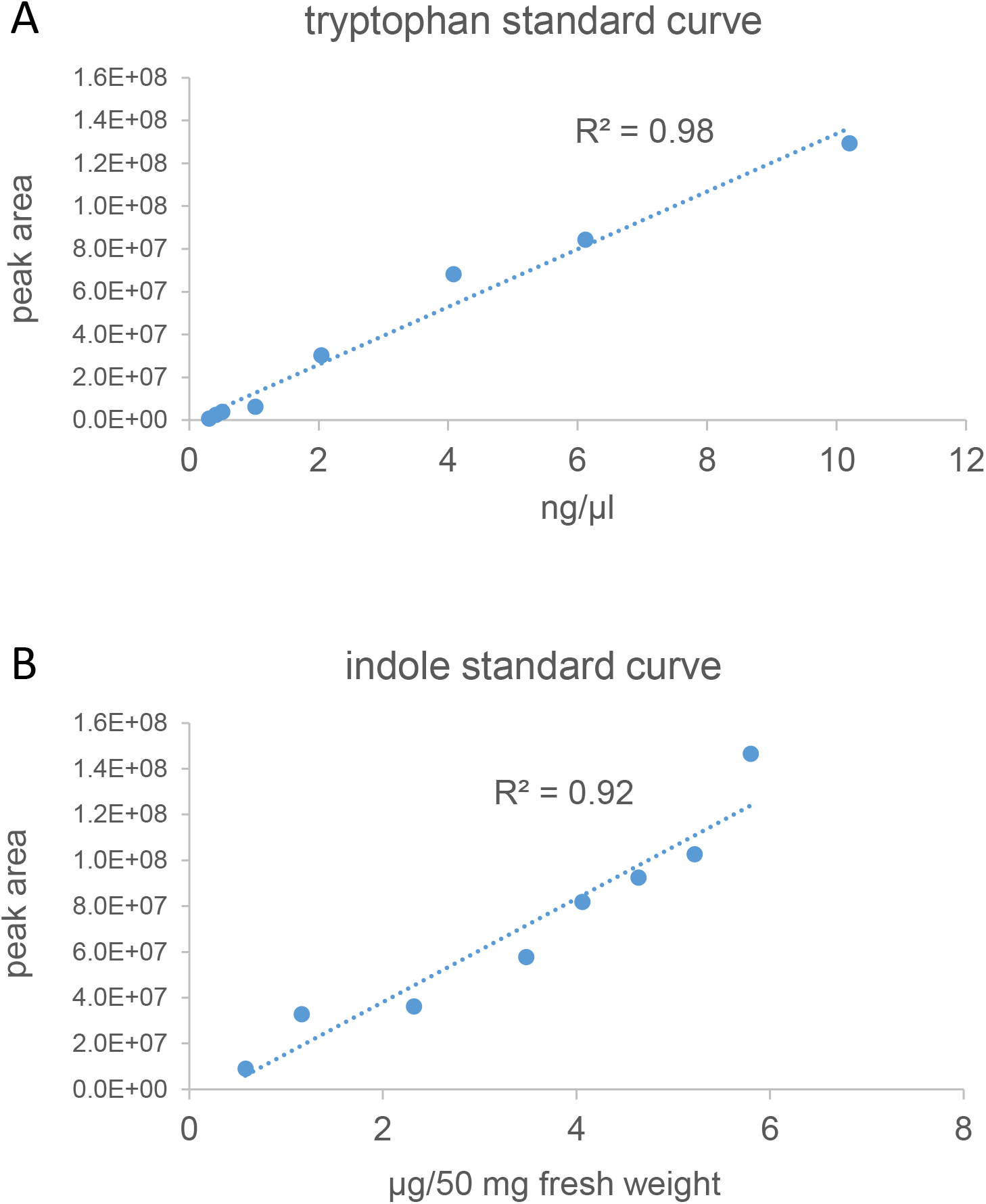
Standard curves for quantifying tryptophan and indole. (A) Tryptophan, as measured by LC-MS, (B) Indole, as measured by GC-MS.

